# Predicting the zoonotic capacity of mammals to transmit SARS-CoV-2

**DOI:** 10.1101/2021.02.18.431844

**Authors:** Ilya R. Fischhoff, Adrian A. Castellanos, João P.G.L.M. Rodrigues, Arvind Varsani, Barbara A. Han

**Author notes:** corresponding author, Barbara A. Han. contributed equally (Ilya R. Fischhoff, Adrian A. Castellanos). **Email addresses:** Ilya Fischhoff, Adrian A. Castellanos, João P.G.L.M. Rodrigues, Arvind Varsani.

## Abstract

Back and forth transmission of SARS-CoV-2 between humans and animals may lead to wild reservoirs of virus that can endanger efforts toward long-term control of COVID-19 in people, and protecting vulnerable animal populations that are particularly susceptible to lethal disease. Predicting high risk host species is key to targeting field surveillance and lab experiments that validate host zoonotic potential. A major bottleneck to predicting animal hosts is the small number of species with available molecular information about the structure of ACE2, a key cellular receptor required for viral cell entry. We overcome this bottleneck by combining species’ ecological and biological traits with 3D modeling of virus and host cell protein interactions using machine learning methods. This approach enables predictions about the zoonotic capacity of SARS-CoV-2 for over 5,000 mammals — an order of magnitude more species than previously possible. The high accuracy predictions achieved by this approach are strongly corroborated by *in vivo* empirical studies. We identify numerous common mammal species whose predicted zoonotic capacity and close proximity to humans may further enhance the risk of spillover and spillback transmission of SARS-CoV-2. Our results reveal high priority areas of geographic overlap between global COVID-19 hotspots and potential new mammal hosts of SARS-CoV-2. With molecular sequence data available for only a small fraction of potential host species, predictive modeling integrating data across multiple biological scales offers a conceptual advance that may expand our predictive capacity for zoonotic viruses with similarly unknown and potentially broad host ranges.

## Introduction

The ongoing COVID-19 pandemic has surpassed 3.9 million deaths globally as of 25 June 2021 [1,2]. Like previous pandemics in recorded history, COVID-19 originated from the spillover of a zoonotic pathogen, SARS-CoV-2, a betacoronavirus originating from an unknown animal host [3–6]. The broad host range of SARS-CoV-2 is due in part to its use of a highly conserved cell surface receptor to enter host cells, the angiotensin-converting enzyme 2 receptor (ACE2) [7] found in all major vertebrate groups [8].

The ubiquity of ACE2 coupled with the high prevalence of SARS-CoV-2 in the global human population explains multiple observed *spillback* infections since the emergence of SARS-CoV-2 in 2019 (see natural infections listed in Table 1 with references). In spillback infection, human hosts transmit SARS-CoV-2 virus to cause infection in non-human animals. In addition to threatening wildlife and domestic animals, repeated spillback infections may lead to the establishment of new animal hosts from which SARS-CoV-2 can continue to pose a risk of *secondary spillover* infection to humans through bridge hosts (e.g., [9]) or newly established enzootic reservoirs. Indeed, this risk has already been realized in Denmark [10] and The Netherlands, where SARS-CoV-2 spilled back from humans to farmed mink (*Neovison vison*) with secondary spillover of a SARS-CoV-2 variant from mink back to humans [11]. A major concern in such secondary spillover events is the appearance of a mutant strain [11,12] affecting host range [13] or leading to increased transmissibility in humans [14,15] (but see [16,17]). Preliminary evidence shows that the mink-derived variant exhibits moderately reduced sensitivity to neutralizing antibodies [10], raising concerns that humans may eventually experience infections from spillback variants, and that vaccines may become less efficient at conferring immunity to these variants [18]. Conversely, human-derived variants pose spillback risks to animals. For example, in contrast to previous infection trials [19], two new human variants are now confirmed to have overcome the species barrier to infect lab mice (*Mus musculus*) [20].

**Table 1.** Species with confirmed suitability for SARS-CoV-2 infection from natural infections or *in vivo* experiments. Asterisks reference species with infection status from preprints (not yet peer-reviewed). Some species (e.g, dogs) with natural infection studies also have *in vivo* experimental studies.

| Species | Susceptibility | Study type | Location | References |
| --- | --- | --- | --- | --- |
| Cow<br>(*Bos taurus*) | Yes | *In vivo*<br>experiment | Lab | [33] |
| Dog<br>(*Canis lupus familiaris*) | Yes | Natural infection | Multiple countries | [34–38] |
| African green monkey<br>(*Chlorocebus aethiops*) | Yes | *In vivo*<br>experiment | Lab | [39] |
| Big brown bat<br>(*Eptesicus fuscus*) | No | *In vivo*<br>experiment | Lab | [40] |
| Cat<br>(*Felis catus*) | Yes | Natural infection | Multiple countries | [35,36,38,41] |
| Gorilla<br>(*Gorilla gorilla*) | Yes | Natural infection | USA, Zoo | [42] |
| Crab-eating macaque<br>(*Macaca fascicularis*) | Yes | *In vivo*<br>experiment | Lab | [43] |
| Rhesus macaque (*Macaca mulatta*) | Yes | *In vivo*<br>experiment | Lab | [44] |
| Golden hamster<br>(*Mesocricetus auratus*) | Yes | *In vivo*<br>experiment | Lab | [45] |
| House mouse<br>(*Mus musculus*) | No | *In vivo*<br>experiment | Lab | [19] (but see [20]) |
| Ferret<br>(*Mustela putorius furo*) | Yes | *In vivo*<br>experiment | Lab | [37] |
| American mink<br>(*Neovison vison*) | Yes | Natural infection | Multiple countries | [35,36,46] |
| Raccoon dog<br>(*Nyctereutes procyonoides*) | Yes | *In vivo*<br>experiment | Lab | [47] |
| European rabbit<br>(*Oryctolagus cuniculus*) | Yes | *In vivo*<br>experiment | Lab | [48] |
| Lion<br>(*Panthera leo*) | Yes | Natural infection | Multiple countries, Zoos | [36,49] |
| Tiger<br>(*Panthera tigris*) | Yes | Natural infection | USA and Sweden, Zoos | [35,36,49,50] |
| Deer mouse<br>(*Peromyscus maniculatus*)* | Yes | *In vivo*<br>experiment | Lab | [51,52] |
| Cougar<br>(*Puma concolor*) | Yes | Natural infection | South Africa, Zoo | [36] |
| Egyptian fruit bat<br>(*Rousettus aegyptiacus*) | Yes | *In vivo*<br>experiment | Lab | [53] |
| Pig<br>(*Sus scrofa*) | No | *In vivo*<br>experiment | Lab | [37,53] |
| Northern treeshrew<br>(*Tupaia belangeri*) | Yes | *In vivo*<br>experiment | Lab | [54] |
| Snow leopard<br>(*Uncia uncia*) | Yes | Natural infection | USA, Zoo | [55] |
| Bank vole<br>(*Clethrionomys glareolus*) | Yes | *In vivo*<br>experiment | Lab | [56] |
| Asian small-clawed otter<br>(*Aonyx cinereus*) | Yes | Natural infection | USA, Zoo | [36,57] |
| White-tailed deer<br>(*Odocoileus virginianus*) | Yes | *In vivo*<br>experiment | Lab | [58] |

Spillback infections from humans to animals are already occurring worldwide. A variety of pets, domesticated animals, zoo animals, and wildlife have also been documented as new hosts of SARS-CoV-2 (Table 1). In addition to secondary spillover infections from mink farms, SARS-CoV-2 has been found for the first time in wild and escaped mink in multiple states in the United States, with viral sequences identical to SARS-CoV-2 in nearby farmed mink [21–23]. The global scale of human infections and the increasing range of known hosts observed for SARS-CoV-2 demonstrate that SARS-CoV-2 has the capacity to establish novel enzootic infection cycles in animals. In response, recent computational studies make predictions about the susceptibility of particular animal species to SARS-CoV-2 [13,24–32]. These studies compare known sequences of ACE2 orthologs across species (*sequence-based* studies), or model the structure of the viral spike protein bound to ACE2 orthologs (*structure-based* studies). These studies yield a wide range of predictions with varying degrees of agreement with laboratory animal experiments (Figure 1).

**Figure 1.**
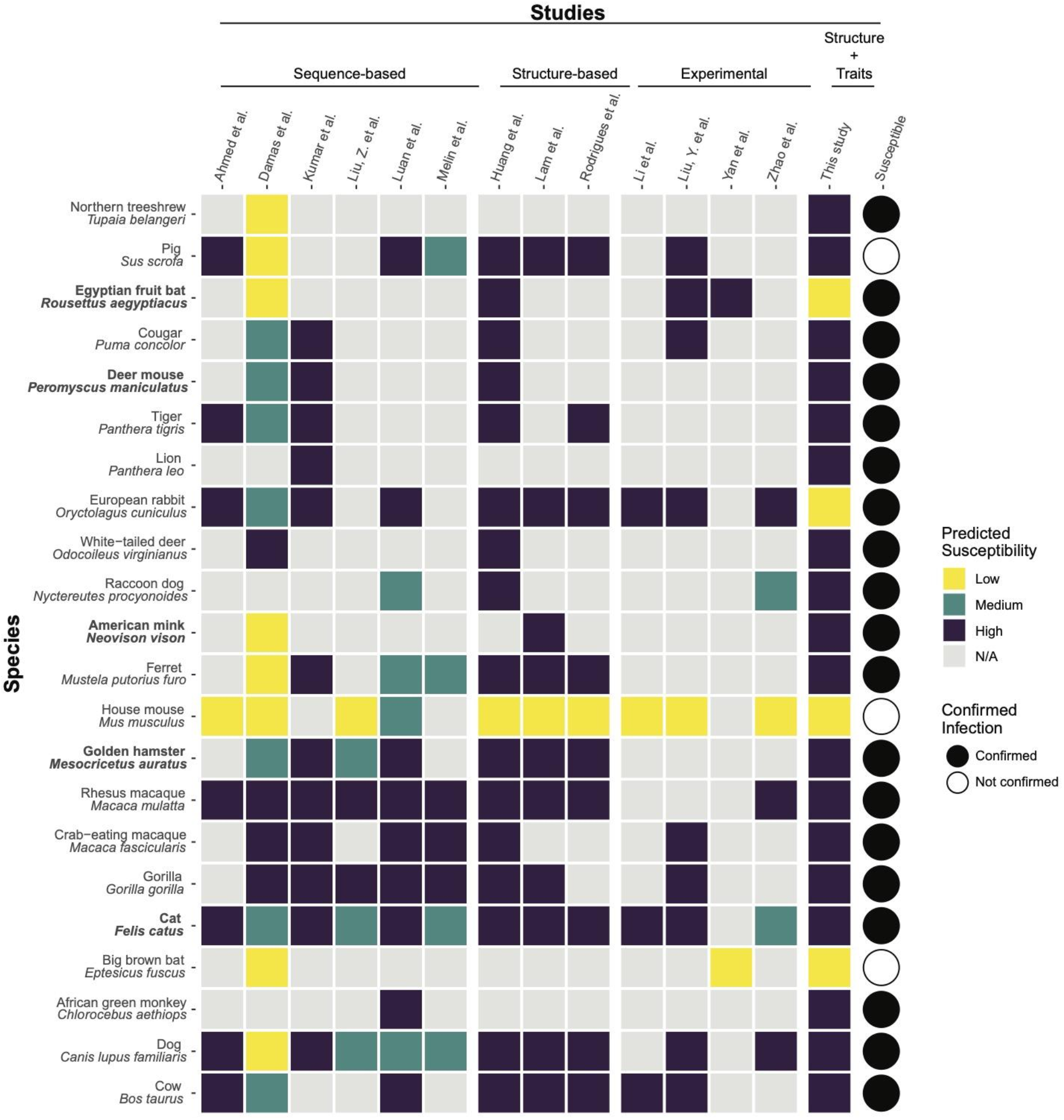
A heatmap summarizing predicted susceptibility to SARS-CoV-2 for species with confirmed infection from *in vivo* experimental studies or from documented natural infections. Studies that make predictions about species susceptibility are shown on the x-axis, organized by method of prediction (those relying on ACE2 sequences, estimating binding strength using three dimensional structures, or laboratory experiments). Predictions about zoonotic capacity from this study are listed in the second to last column, with high and low categories determined by zoonotic capacity observed in *Felis catus*. Confirmed infections for species along the y-axis are summarized in [59] and are depicted as a series of filled or unfilled circles. Bolded species have been experimentally confirmed to transmit SARS-CoV-2 to naive conspecifics. Species predictions range from warmer colors (yellow: low susceptibility or zoonotic capacity for SARS-CoV-2) to cooler colors (purple: high susceptibility or zoonotic capacity). See Supplementary Methods (https://doi.org/10.25390/caryinstitute.c.5293339) for detailed methods about study categorization.

### Sequence-based studies

Sequence-based studies predict host susceptibility based on amino acid sequence similarity between human (hACE2) and non-human ACE2, and assume that a high degree of similarity correlates with stronger viral binding, especially at amino acid residues where hACE2 interacts with the SARS-CoV-2 spike glycoprotein. For some species, such as rhesus macaques [60], these qualitative predictions are borne out by *in vivo* studies (Figure 1), but predictions from these methods do not consistently match real-world outcomes. For example, sequence similarity predicted weak viral binding for minks and ferrets, which have all been confirmed as highly susceptible, with minks capable of onward transmission [11,32,37] (Figure 1). These mismatches to experimental or real-world outcomes may arise in part because protein three-dimensional structure, the main determinant of protein function, is incompletely represented by 1D amino acid sequences [61,62]. As such, details about the interaction between host ACE2 and the viral spike protein are not well captured by sequence-based studies.

### Structure-based studies

Modeling the three-dimensional structure of protein-protein complexes addresses some of the limitations of sequence-based approaches, and has proven useful to predict how different ACE2 orthologs bind to the SARS-CoV-2 viral spike protein receptor-binding domain (RBD) [13,24]. These studies have also identified particular ACE2 amino acid residues essential for a productive interaction with the viral RBD, thus improving predictive models of susceptibility through structure-based inference [13]. These studies leverage known structures of the hACE2 receptor bound to the SARS-CoV-2 RBD and use powerful simulation methods to predict how variation across different ACE2 orthologs affects binding with the viral RBD. While these approaches successfully predicted strong binding for species that have been infected (e.g. domestic cat, tiger, dog, and ferret) and weak binding for species in which experimental infections failed (e.g. chicken, duck [37], mouse [19]), the results are also not consistently supported by experiments. For instance, while guinea pig ACE2 scored favorably among susceptible species in one of the studies [13], this ortholog was shown experimentally not to bind to the SARS-CoV-2 RBD [63].

Although structural modeling has produced the most accurate results to date, all currently available approaches for predicting the host range of SARS-CoV-2 are fundamentally constrained by the availability and quality of ACE2 sequences across species. ACE2 is ubiquitous across chordates, likely because of its role in several highly conserved physiological pathways [64]. Because it is so highly conserved, the vast majority of mammal species (>6,000 species) are likely to have ACE2 receptors, but there are many fewer sequences available from which to make predictions using existing modeling methods (∼300 species). The functional importance of the ACE2 receptor suggests that it has evolved in association with other intrinsic organismal traits that are more easily observed and for which data are available for many more species. These suites of correlated organismal traits may provide a robust statistical proxy that can be leveraged to predict suitable hosts for SARS-CoV-2. Previous trait-based analyses applied statistical (machine) learning techniques to accurately distinguish the zoonotic capacity of various organisms [65–67], and predict likely hosts for particular groups of related viruses [68,69], predictions which have subsequently been validated through independent laboratory and field investigations (e.g., [70,71]).

Here, we combine molecular structural modeling of viral binding with machine learning of species-level ecological and biological traits to predict species’ zoonotic capacity for SARS-CoV-2 virus across 5,400 mammal species, expanding our predictive capacity by an order of magnitude (Figure 2). Crucially, this integrated approach enables predictions for the vast majority of species whose ACE2 sequences are currently unavailable by leveraging information from viral binding dynamics and biological traits of potential hosts. In our workflow (Figure 2), we first carry out structural modeling to quantify the binding strength of SARS-CoV-2 RBD for vertebrate species using published ACE2 amino acid sequences [72]. We then collate species traits and train a machine learning model to predict the zoonotic capacity for 5,400 mammal species. Zoonotic capacity (host susceptibility and capacity for onward transmission) was approximated through a conservative threshold of binding strength applied to our structural modeling results and reported by *in vivo* studies.

**Figure 2.**
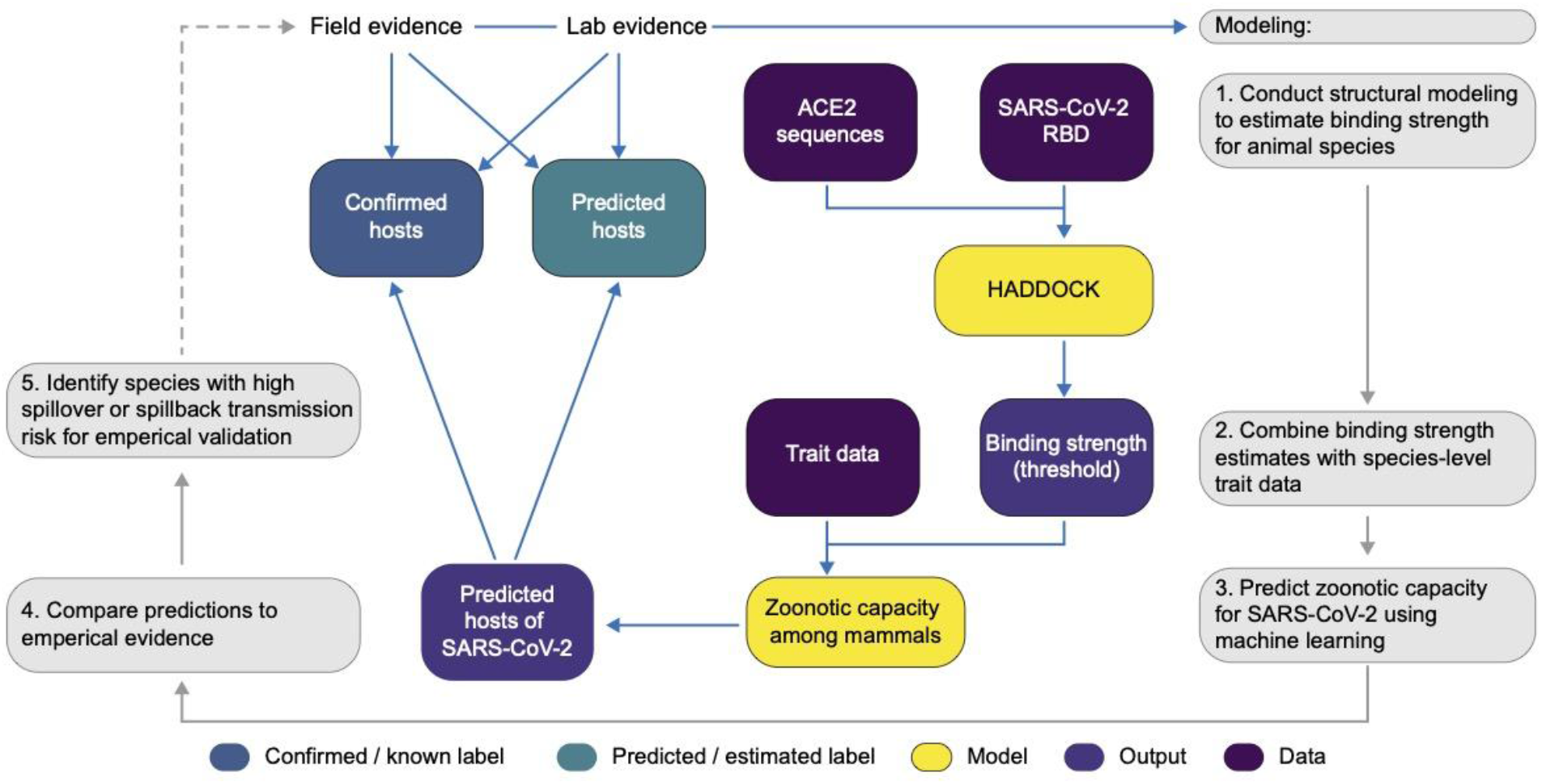
A flowchart showing the progression of our workflow combining evidence from limited lab and field studies with additional data types to predict zoonotic capacity across mammals through multi-scale statistical modeling (gray boxes, steps 1-5). For all vertebrates with published ACE2 sequences, we modelled the interface of species’ ACE2 bound to the viral receptor binding domain using HADDOCK. We then combined the HADDOCK scores, which approximate binding strength, with species’ trait data and trained machine learning models for both mammals and vertebrates (yellow boxes). Mammal species predicted to have high zoonotic capacity were then compared to results of *in vivo* experiments and *in silico* studies that applied various computational approaches. Based on predictions from our model, we identified a subset of species with particularly high risk of spillback and secondary spillover potential to prioritize additional lab validation and field surveillance (dashed line).

COVID-19 is, at this time, primarily a disease affecting humans, thus spillback infection of SARS-CoV-2 from humans to animals is the most likely mode by which new host species will become established. We therefore identify a subset of species for which the threat of spillback infection appears greatest due to geographic overlaps and opportunities for contact with humans in areas of high SARS-CoV-2 prevalence globally. Our predictions contribute to a critical interdisciplinary and iterative process between computational modeling, field surveillance, and laboratory experiments that is necessary for improving zoonotic risk quantification, and to better inform next steps toward the prevention of enzootic SARS-CoV-2 transmission and spread. We demonstrate our approach using the SARS-CoV-2 sequence that initially emerged in humans. These methods can be readily expanded to enable host range predictions for new variants as their hACE2-RBD crystal structures become available.

## Methods

### Protein sequence and alignment

We assembled a dataset of ACE2 NCBI GenBank accessions that are known human ACE2 orthologs or have high similarity to known orthologs as determined using BLASTx [73]. Using the R package *rentrez* and the accession numbers, we downloaded ACE2 protein sequences [74]. We supplemented these sequences by manually downloading four additional sequences from the MEROPS database [75].

### Structural Modeling of ACE2 orthologs bound to SARS-CoV-2 spike

The modeling of all 326 ACE2 orthologs bound to SARS-CoV-2 spike receptor binding domain was carried out as described previously [13], with a few differences. Sequences of ACE2 orthologs were aligned using MAFFT [76] and trimmed to the region resolved in the template crystal structure of hACE2 bound to the SARS-CoV-2 spike (PDB ID: 6m0j, [77]. Ambiguous positions in each sequence, artifacts of the sequencing method, were replaced by Glycine to minimize assumptions about the nature of the amino acid side-chain but still allow for modeling. For each ortholog, we generated 10 homology models using MODELLER 9.24 [78,79], with restricted optimization (*fastest* schedule) and refinement (*very_fast* schedule) settings, and selected a representative model based on the normalized DOPE score. These representative models were then manually inspected and 27 were removed from further analysis due to large insertions/deletions or to the presence of too many ambiguous amino acids at the interface with spike. Each validated model was submitted for refinement to the HADDOCK web server [80], which ran 50 independent short molecular dynamics simulations in explicit solvent to optimize the interface between the two proteins. For each one of the animal species in our study, we assigned an average and standard deviation of the scores of the 10 best refined models, ranked by their HADDOCK score -- a combination of van der Waals, electrostatics, and desolvation energies. A lower (more negative) HADDOCK score predicts stronger binding between the two proteins. We hereafter refer to predicted binding strength, or simply binding strength, to indicate HADDOCK score. The HADDOCK server is freely available, and we provide code to reproduce analyses or to aid in the application of this modeling approach to other similar problems (https://zenodo.org/record/4517509).

### Trait data collection and cleaning

We gathered ecological and life history trait data from AnAge [81], Amniote Life History Database [82], and EltonTraits [83], among other databases (Supplementary Table 1; for details on data processing, see Supplementary Methods with all supplementary data, figures, methods, and tables available at https://doi.org/10.25390/caryinstitute.c.5293339). Using these data, we also engineered additional traits that have shown importance in predicting host-pathogen associations in other contexts. For example, as a measure of habitat breadth [84], we computed for each species the percentage of ecoregions it occupies. To assess the influence of sampling bias across species, we used the wosr R package [85] to count the number of studies returned in a search in Web of Science for each species’ Latin binomial and included this as a proxy for sampling bias in our model.

Following the results of initial structural modeling (described above), we observed that per-residue energy decomposition analysis of HADDOCK scores for 29 species indicated that all species with strong predicted binding had in common a salt bridge between SARS-CoV-2 K417 and a negatively charged amino acid at position 30 in the ACE2 sequence [13]. Given the apparent effect of amino acid 30 on overall binding strength, we constructed an additional feature to denote whether amino acid 30 is negatively charged (and therefore more likely to support strong binding) and included this feature as an additional trait in our models.

### Modeling

#### Quantifying a threshold for zoonotic capacity using HADDOCK

While ACE2 binding is necessary for viral entry into host cells, it is not sufficient for SARS-CoV-2 transmission. Multiple *in vivo* experiments suggest that not all species that are capable of binding SARS-CoV-2 are capable of transmitting active infection to other individuals (e.g., cattle, *Bos taurus* [33]; bank voles, *Myodes glareolus [56]*). Viral replication, and infectious viral shedding that enables onward transmission, are both required for a species to become a suitable bridge or reservoir species for SARS-CoV-2. In order to constrain our predictions to species with the greatest potential to perpetuate onward transmission, we trained our models on a conservative threshold of binding strength (HADDOCK score = -129). This value is between the scores for two species: the domestic cat (*Felis catus*), which is currently the species with weakest predicted binding with confirmed conspecific transmission [86], and the pig (*Sus scrofa*), which shows the strongest estimated binding for which experimental inoculation failed to cause detectable infection [37]. Binding strength was binarized according to this threshold, above which it is more likely that both infection and onward transmission will occur following the results of multiple empirical studies (Table 1). We note that there are species confirmed to be susceptible whose predicted binding strength is weaker than cats, but conspecific transmission has not been confirmed in these species. While it is likely that intraspecific transmission will be reported for additional species as the pandemic continues, the binding strength selected for this analysis represents an appropriately conservative threshold based on currently available evidence. For additional modeling details, see Supplementary Methods.

### Trait-based modeling to predict zoonotic capacity

We applied generalized boosted regression [87] to host trait data to predict species’ binding strength to SARS-CoV-2. We applied this approach initially to all of the vertebrate species for which we estimated HADDOCK scores, but these models did not perform well. This was likely due to extensive dissimilarities among traits describing different classes of organisms. For instance, traits that are commonly measured for reptiles are different from those of interest for birds or amphibians. Moreover, currently available ACE2 sequences are dominated by ray-finned fishes and mammals.

Given that only mammals have so far been confirmed as both susceptible and capable of onward transmission of SARS-CoV-2, we created a separate set of models to make zoonotic capacity predictions for mammals only. For this mammal-only dataset, we gathered additional species-level traits from PanTHERIA [88] and added a series of binary fields for taxonomic order (based on [89]; Supplementary Table 2). We then applied boosted regression (BRT; gbm package [90] in R version 4.0.0 ^[90,91]^) to impute missing trait data for mammal species (e.g., [67]; see Supplementary Methods for details on imputation methods and results).

Many of the mammals for which we found the strongest evidence of zoonotic capacity are domesticated to some degree (pets, farmed or traded animals, lab models) [11,37,53]. Relative to their ancestors or wild conspecifics, domesticated animals often have distinctive traits [92] that are likely to influence the number of zoonoses found in these species [93]. To account for trait variation due to domestication in certain species, we modeled mammals in two ways. First, we incorporated a variable indicating whether the source populations from which trait data were collected are wild or non-wild (e.g., farmed, pets, laboratory animals; non-wild status confirmed by the Mammal Diversity Database [94]). Trait data collected from both wild and non-wild individuals were considered to represent non-wild species for the purposes of this model. In a second approach, we used only the wild species for model training and evaluation. For both approaches, pre-imputation trait values were used for all non-wild mammals during model training, evaluation, and prediction.

Boosted regression (BRT) is an ensemble machine learning approach that accommodates non-random patterns of missing data, nonlinear relationships, and interacting effects among predictors. In a BRT model, a sequence of regression models are fit by recursive binary splits, with each additional regression modeling those instances that were poorly accounted for by the previous regression iterations in the tree [87]. We applied grid search to select optimal hyperparameters, and repeated model fitting 50 times using bootstrapped training sets of 80% of labeled data. We measured performance by the area under the receiver operating characteristic curve (AUC) for predictions made on the test dataset (remaining 20%), corrected by comparing with null models created by target shuffling, which employed similar bootstrapping (50 times). Detailed methods can be found in Supplementary Methods. We discuss herein the results of model predictions about zoonotic capacity made by applying this final model to all mammal species. We also report the mean and variation in predicted probabilities across all 50 bootstrapped models in Supplementary File 1.

To visualize geographic patterns, we mapped the geographic ranges of mammal species predicted within the 90th percentile of zoonotic capacity for SARS-CoV-2 using International Union for the Conservation of Nature (IUCN) polygons of species distributions [95]. We subset to the species found in human-associated habitats (e.g., urban areas, crop lands, heavily degraded forests; based on IUCN 2020), and also masked their ranges to areas of high human case counts (using SARS-CoV-2 case data from the COVID-19 Data Repository at Johns Hopkins University [1]).

Additional methods and results of other uninformative model variations are also described in Supplementary Methods and Supplementary Table 3 (e.g., a model in which binding strength is modeled as a continuous rather than a threshold measure, a model predicting the charge at amino acid 30, a model for all vertebrate species) (https://doi.org/10.25390/caryinstitute.c.5293339). We provide code and data files for carrying out boosted regression tree models (https://github.com/HanLabDiseaseEcology/zoonotic_capacity). Details about how the species susceptibility predictions from past studies were standardized into categories (low, medium, high; Figure 1) are also available in Supplementary Methods.

## Results

### ACE2 host protein sequences and alignment

The ACE2 protein sequence alignment of the orthologs from 326 species spans eight classes and 87 orders (https://zenodo.org/record/4517509). The majority of sequences belonged to the classes Actinopterygii (22.1%), Aves (23.3%), and Mammalia (46.6%). Sequence length ranged from 344 amino acids to 872 with a median length of 805.

### Structural modeling of viral binding strength

We predicted binding strength for 299 vertebrates, including 142 mammals. These binding strength scores represented six classes and 80 orders and ranged between -167.816 and -105.615. Across these six vertebrate classes, the strongest predicted binding between ACE2 and SARS-CoV-2 (corresponding to the lowest mean HADDOCK scores) were in ray-finned fishes (Actinopterygii; mean = -137.945) and mammals (Mammalia; mean = -129.193) (Figure 3A). Four of these six classes included at least one species predicted to have stronger binding than *Felis catus* (Figure 3B). Among well-represented mammalian orders (those containing at least 10 species with binding strength predictions), Primates and Carnivora showed predicted mean binding strengths that were stronger than domestic cats (Figure 3C).

**Figure 3.**
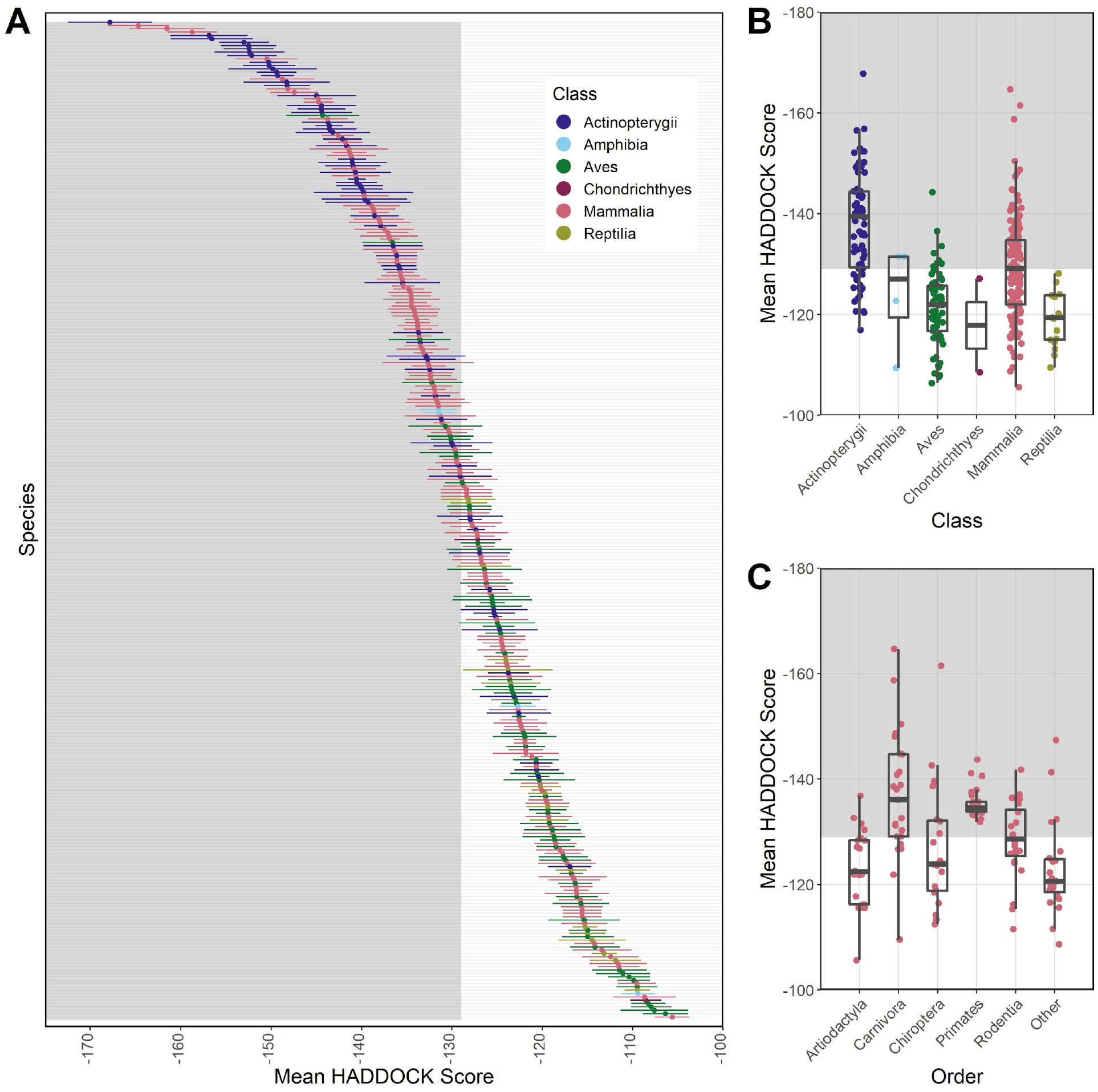
Plots showing results from modeling species’ ACE2 interaction with SARS-CoV-2 RBD using HADDOCK to predict binding strength (measured as arbitrary units). HADDOCK scores that predict stronger binding are more negative. The mean and standard deviation of the HADDOCK score for vertebrate species **(A)** for which ACE2 orthologs are available. Binding strengths vary across vertebrate classes **(B)** and across the five most speciose mammalian orders **(C)**. The “Other” category contains species across multiple orders for which ACE2 sequences were available, each with fewer than 10 representative species in the order. The shaded regions of all panels represent predicted binding that is as strong or stronger than (more negative values than) the domestic cat (*Felis catus*), which represents our conservative zoonotic capacity threshold based on currently available empirical evidence.

### Species predictions of zoonotic capacity from trait-based machine learning models

The best performing model was trained on a mammal-only dataset with trait imputation and showed corrected test AUC of 0.72 (for results of all other model variations, see Supplementary Table 3). We used this model to generate predictions of zoonotic capacity among mammal species. Citation count, as a proxy for study effort, had ∼1% relative importance, suggesting that sampling bias across species had little influence on the model.

This zoonotic capacity model identified 540 species (out of 5400 total mammal species) within the 90th percentile probability (0.826 or higher, compared to a total of 2,401 mammal species with prediction scores above 0.5; see Supplementary File 1 for predictions on all 5,400 species, https://doi.org/10.25390/caryinstitute.c.5293339). The top 10% of species with the highest predicted probabilities includes representatives from 13 orders. Most primates were predicted to have high zoonotic capacity and collectively showed stronger viral binding compared to other mammal groups (Figure 4). Additional orders with numerous species predicted to have high zoonotic capacity (at least 75% of species above 0.5) include Hyracoidea (hyraxes), Perissodactyla (odd-toed ungulates), Scandentia (treeshrews), Pilosa (sloths and anteaters), Pholidota (pangolins), and non-cetacean Artiodactyla (even-toed ungulates) (Figure 4).

**Figure 4.**
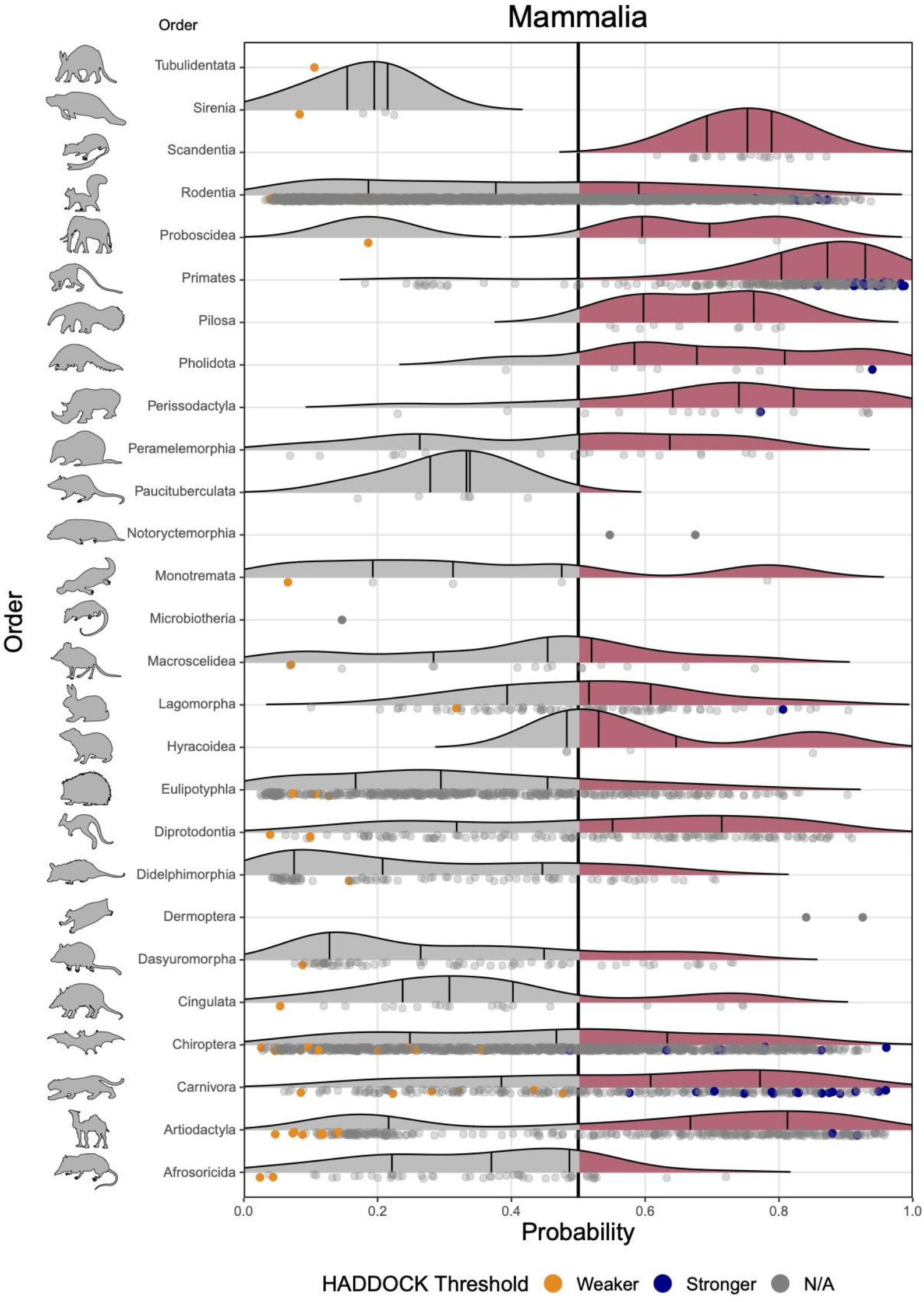
Ridgeline plots showing the distribution of predicted zoonotic capacity across mammals. Predicted probabilities for zoonotic capacity across the x-axis range from 0 (likely not susceptible) to 1 (zoonotic capacity predicted to be the same or greater than *Felis catus*), with the vertical line representing 0.5. The y-axis depicts all mammalian orders represented by our predictions. Density curves represent the distribution of the predictions, with those parts of the curve over 0.5 colored pink and lines representing distribution quartiles. The predicted values for each order are shown as points below the density curves. Points that were used to train the model are colored: orange represents species with weaker predicted binding, blue represents species with stronger predicted binding. Selected family-level distributions are shown in the Supplemental Figures 5-6 (https://doi.org/10.25390/caryinstitute.c.5293339).

### Comparison of species predictions

#### Comparing species predictions across multiple computational approaches

Our model combined species traits with estimates of viral binding strength to predict zoonotic capacity, which encompasses both susceptibility to SARS-CoV-2 and the probability of onward transmission. Zoonotic capacity was defined as a threshold value based on the results of experimental studies confirming intraspecific transmission among animals, and is therefore more conservative than thresholds adopted by other studies (e.g., those based only on estimates of viral binding strength, [30]). In addition, our modeling approach (machine learning) and prediction targets (zoonotic capacity) differed compared to existing computational approaches, which applied sequence-based or structure-based analyses constrained by the small number of published ACE2 sequences. Despite these differences, comparing the species predictions generated by multiple different approaches can be useful for gauging consensus, and for comparing how species predictions change from one method to another.

Across approaches, there was general agreement in the predictions for primates as well as for a select group of artiodactyls and carnivores (Figure 5). Our model results also agreed with low susceptibility predictions made by several previous studies using sequence-based approaches (e.g., in certain bats and rodents). In general, we note that structure-based models predicted a smaller proportion of species to have low susceptibility compared to sequence-based studies.

**Figure 5.**
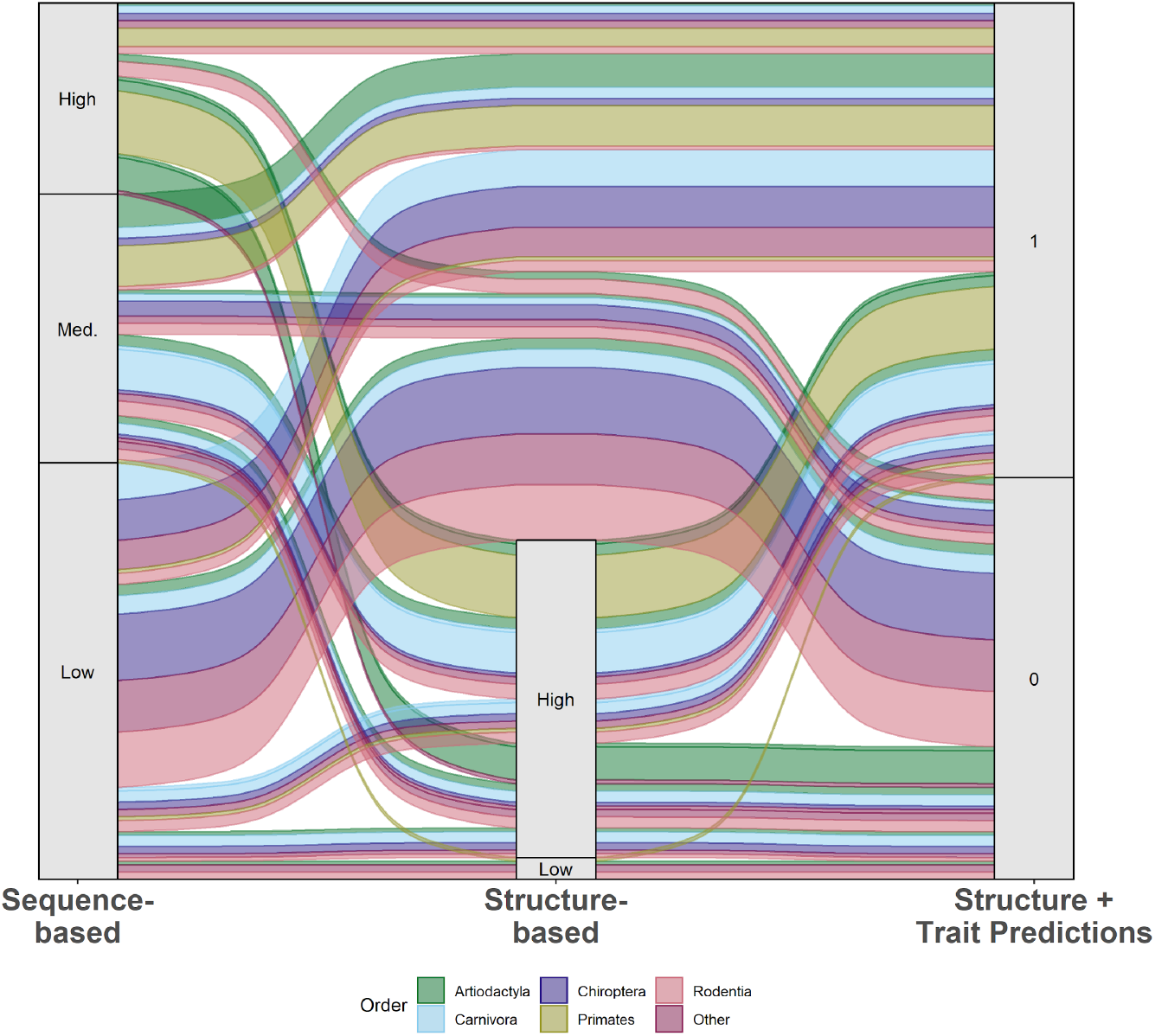
An alluvial plot comparing predictions of species susceptibility from multiple methods. Existing studies (listed in Supplementary Methods) are categorized as either sequence-based or structure-based. Predictions from our zoonotic capacity model result from combining structure-based modeling of viral binding with organismal traits using machine learning to distinguish species with zoonotic capacity above (1) or below (0) a conservative threshold value set by domestic cats (*Felis catus*). Colors represent unique mammalian orders, and the width of colored bands represent the relative number of species with that combination of predictions across methods. See Supplementary Methods (https://doi.org/10.25390/caryinstitute.c.5293339) for details on how species across multiple studies were assigned to categories (high, medium, low).

#### Comparing model predictions to *in vivo* outcomes

Our model predictions matched the results of several recently published *in vivo* studies on SARS-CoV-2 infection (Figure 1). For instance, experiments on deer mice (*Peromyscus maniculatus*; [51,52]) and raccoon dogs (*Nyctereutes procyonoides*; [47]) confirmed SARS-CoV-2 infection and transmission to naive conspecifics. Our model also estimated a high probability of zoonotic capacity of American mink for SARS-CoV-2 (*Neovison vison*, probability=0.83, 90th percentile), in which farmed individuals present severe infection from human spillback, and demonstrate the capacity to transmit to conspecifics as well as to humans [11,46]. Our model also correctly predicted relatively low zoonotic capacity for big brown bats (*Eptesicus fuscus*; [40]).

There were notable differences between our model results and the outcomes of some experimental studies. For instance, our model estimated a moderately high probability of zoonotic capacity for pigs (*Sus scrofa*, probability = 0.72, ∼80th percentile). Similarly, some computational and cell-based studies have also predicted strong viral binding in this species [26,96], but *in vivo* studies report no detectable infection or onward transmission of SARS-CoV-2 [37,53]. Similarly for cattle (*Bos taurus*), our model estimated a moderately high probability for zoonotic capacity (0.72, ∼80th percentile), and in a live animal experiment, cattle were confirmed to be susceptible to infection but no onward transmission was observed to virus-naive conspecifics [33].

## Discussion

We combined structure-based models of viral binding with species-level data on biological and ecological traits to make predictions about the capacity of animal species to become zoonotic hosts of SARS-CoV-2 (*zoonotic capacity*). This combined modeling approach predicted zoonotic capacity with 72% accuracy, extending our predictive capacity beyond the limited number of species for which ACE2 sequences are currently available. We identified numerous mammal species whose predicted zoonotic capacity meets or exceeds the viral susceptibility and transmissibility observed in experimental infections with SARS-CoV-2. In addition to wide agreement with *in vivo* study results produced to date (Table 1), these model predictions corroborate the predictions of previous studies generated using the limited number of available ACE2 sequences (Figure 1). Below we discuss predictions of zoonotic capacity for a number of ecologically and epidemiologically relevant categories of mammalian hosts.

### Captive, farmed, or domesticated species

Given that the type and frequency of contact with humans fundamentally underlies transmission risk, it is notable that our model predicted high zoonotic capacity for multiple captive species that have also been confirmed as susceptible to SARS-CoV-2 via experiments or natural infections. These include numerous carnivore species, such as large cats from multiple zoos and pet dogs and cats. Our model also predicted high SARS-CoV-2 zoonotic capacity for many farmed, domesticated, and live traded species. The water buffalo (*Bubalus bubalis*), widely bred for dairy production and farming, had the highest probability of zoonotic capacity among livestock (0.91). Model predictions in the 90th percentile also included American mink (*Neovison vison*), red fox (*Vulpes vulpes*), sika deer (*Cervus nippon*), white-lipped peccary (*Tayassu pecari*), nilgai (*Boselaphus tragocamelus*), and raccoon dogs (*Nyctereutes procyonoides*), all of which are farmed, with the latter two considered invasive species in some areas [97,98]. In addition to the risks of secondary spillover to humans and the potential for large economic losses from culling infected animals [99], the escape of farmed individuals into wild populations has implications for the spread and enzootic establishment of SARS-CoV-2 [21]. These findings also have implications for vaccination strategies, for instance, prioritizing people in regular contact with potential bridge species (e.g., veterinarians, abattoir-workers, farmers, etc).

### Live traded or hunted wildlife species

The majority of the legally traded live mammals are primates and carnivores [100], and model predictions included several species from these groups. Our model predicted high zoonotic capacity in 20 out of 21 species in the primate genus *Macaca*, which comprise the majority of all live-traded primates. Several live-traded carnivores and pangolins were also assigned high zoonotic capacity, including the Asiatic black bear (*Ursus thibetanus*), grey wolf (*Canis lupus*), and jaguar (*Panthera onca*), the Philippine pangolin (*Manis culionensis*) and Sunda pangolin (*M. javanica*). Pangolins are notable because one of the betacoronaviruses with the highest sequence similarity to SARS-CoV-2 was isolated from Sunda pangolins [101,102]. Pangolin burrows are also known to be occupied by multiple other animal species, including numerous bats [103].

Commonly hunted species in the top 10% of predictions include duiker (*Cephalophus zebra*, West Africa), warty pig (*Sus celebes*, Southeast Asia), and two species of deer (*Odocoileus hemionus* and *O. virginianus*) that are widespread across the Americas. The white-tailed deer (*O. virginianus*) was recently confirmed to be capable of transmitting SARS-CoV-2 to conspecifics via indirect contact (aerosolized virus particles) [58].

### Bats

Similarly, bats are of special interest because of the high diversity of betacoronaviruses found in *Rhinolophus spp*. and other bat species [104–107]. Our model identified 35 bat species within the 90th percentile of zoonotic capacity for SARS-CoV-2. Within the genus *Rhinolophus*, our model identified the large rufous horseshoe bat (*Rhinolophus rufus*), a known natural host for bat betacoronaviruses [104] and a congener to three other horseshoe bats harboring betacoronaviruses with high nucleotide sequence similarity to SARS-CoV-2 (∼92-96%) [6,108,109]. For these three species, our model assigned a range of probabilities for SARS-CoV-2 zoonotic capacity (*Rhinolophus affinis* (0.58), *R. malayanus* (0.70), and *R. shameli* (0.71)) and also predicted relatively high probabilities for two congeners, *Rhinolophus acuminatus* (0.84) and *R. macrotis* (0.70). These predictions are in agreement with recent experiments demonstrating efficient viral binding of SARS-CoV-2 RBD for *R. macrotis* [110] and confirmation of SARS-CoV-2-neutralizing antibodies in field-caught *R. acuminatus* harboring a closely related betacoronavirus [111].

Our model also identified 17 species in the genus *Pteropus* (flying foxes) with high probabilities of zoonotic capacity for SARS-CoV-2. Some of these species are confirmed reservoirs of other zoonotic viruses in Southeast Asia (e.g., henipaviruses in *P. lylei, P. vampyrus, P. conspicillatus*, and *P. alecto*). While contact patterns between bats and humans may be somewhat less direct compared with captive or farmed species, annual outbreaks attributed to viral spillover transmission from bats illustrate a persistent epizootic risk to humans [112–114] and confirm that gaps in systematic surveillance of zoonotic viruses, including betacoronaviruses, remain an urgent priority (e.g., [115]).

### Rodents

Our model identified 76 rodent species with high zoonotic capacity for SARS-CoV-2, some of which thrive in human-altered settings. Among these, the deer mouse (*Peromyscus maniculatus*) and the white-footed mouse (*P. leucopus*) showed high probabilities. These are among the most well-studied mammals in North America, in part due to their status as zoonotic reservoirs for multiple zoonotic pathogens and parasites [116–118]. Experimental infection, viral shedding, and sustained intraspecific transmission of SARS-CoV-2 were recently confirmed for *P. maniculatus* [51,52], but similar studies have not been conducted for *P. leucopus*, which is widely distributed across the eastern United States and Mexico.

Our model predicted low zoonotic capacity for *Mus musculus* (0.11), corresponding with *in vivo* experiments suggesting this species is not susceptible to infection by the initial human variant of SARS-CoV-2[19], although notably, more recent experiments have confirmed the susceptibility of *M. musculus* to two newer human-derived variants [20]. Also in the top 10% were two rodent species considered to be human commensals whose geographic ranges are expanding due to human activities: *Rattus argentiventer* (0.84) and *R. tiomanicus* (0.79) (Supplementary File 1) [119–121]. Additional common rodent species with relatively high probabilities of zoonotic capacity include domesticated guinea pigs (*Cavia porcellus*), gerbils (*Gerbillus gerbillus, Meriones tristrami*), and several common mouse species (*Apodemus peninsulae, A. flavicollis*, and *A. sylvaticus*), all of which are known reservoirs for other zoonotic diseases [122–124]. It is notable that many of these rodent species are regularly preyed upon by carnivore species, such as the red fox (*Vulpes vulpes*) or domestic cats (*Felis catus*) who themselves were predicted to have high zoonotic capacity for SARS-CoV-2 by our model.

### Species with large geographic ranges

With sufficient opportunity for infectious contact, the risk of zoonotic spillback transmission increases with SARS-CoV-2 prevalence in human populations. Among species with high model-predicted zoonotic capacity, there were several relatively common species with very large geographic ranges or synanthropic tendencies that overlap with global hotspots of COVID-19 in people (Figure 6, Supplementary File 2). Notable species that are widely distributed across much of the northern hemisphere include the red fox (*Vulpes vulpes*, ∼50 countries), the European polecat (*Mustela putorius*), the raccoon dog (*Nyctereutes procyonoides*), stoat (*Mustela erminea*) and wolf (*Canis lupus*). White-tailed deer (*Odocoileus virginianus*) are among the most geographically widespread species across Latin American countries with high SARS-CoV-2 prevalence. Globally, South and Southeast Asia had the highest diversity of mammal species with high predicted zoonotic capacity for SARS-CoV-2 (∼90 species). Notable examples in this region include both rodents and bats. For example, Finlayson’s squirrel (*Callosciurus finlaysonii*) is native to Mainland Southeast Asia, but introductions via the pet trade in Europe have led to invasive populations in multiple countries [125]. Hunting has been documented for numerous bat species with geographic ranges across Southeast Asia (e.g., *Cheiromeles torquatus, Cynopterus brachyotis, Rousettus amplexicaudatus, Macroglossus minimus*) [126,127], and there were multiple additional bat species in the 90th percentile from Asia and Africa where bats are subject to hunting pressure and from which other betacoronaviruses have been identified [107,128]. There were also several wide-ranging species whose contact with humans are limited to specialized settings. For instance, biologists and wildlife managers handle live individuals for research purposes, including grizzly bear (*Ursus arctos*), polar bear (*Ursus maritimus*), and wolf (*Canis lupus*), all of which are in the 89th percentile or above for predicted zoonotic capacity to SARS-CoV-2.

**Figure 6:**
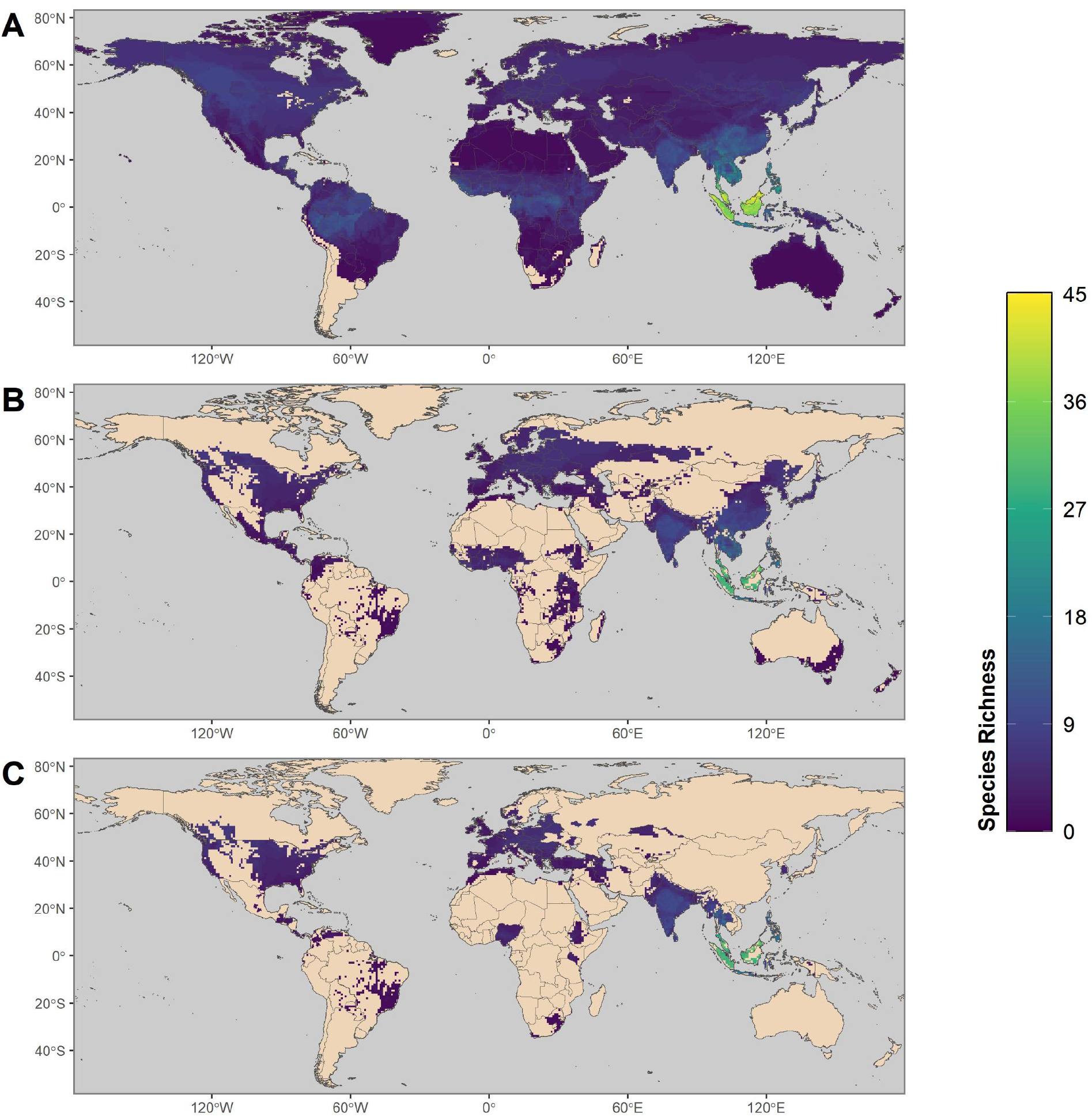
Maps showing the global distribution of species with predicted capacity to transmit SARS-CoV-2. **(A)** depicts global species richness of the top 10 percent of model-predicted zoonotic capacity. Geographic ranges of this subset of species were filtered to those associated with human-dominated or human-altered habitats **(B)**, and further filtered to show the subset of species that overlaps with areas of high human SARS-CoV-2 positive case counts (over 100,000 cumulative cases as of 17 May 2021) **(C)**. For a full list of model-predicted zoonotic capacity of species by country, see Supplementary File 2 (https://doi.org/10.25390/caryinstitute.c.5293339).

### Other high priority mammal species

Species with more equivocal predictions about zoonotic capacity that are in frequent contact with humans warrant further investigation. For instance, while species such as horses (*Equus caballus*), goats (*Capra hircus*), and guinea pigs (*Cavia porcellus*) are not in the top 10% of predicted zoonotic capacity, due to the nature of their contact with humans they may experience greater risks of spillback infection, or pose a greater risk to humans for secondary spillover infection compared to many wild species. Conversely, while certain endangered or nearly extinct species are predicted to have relatively high zoonotic capacity, they may have fewer opportunities for human contact. For species of conservation concern, spillback transmission of SARS-CoV-2 from humans presents an important source of risk [28,129], particularly for populations that are under active management, including *ex situ* management such as captive breeding. These species include the scimitar-horned oryx (*Oryx dammah*), addax (*Addax nasomaculatus*), some Antarctic fauna, and mountain gorillas (*Gorilla beringei*) in which SARS-CoV-2 spillback infection may occur through close-proximity eco-tourism activities [130,131]. Indeed, spillback transmission of SARS-CoV-2 has already been confirmed in a closely related species, the Western lowland gorilla (*Gorilla gorilla*) in captivity [132], leading to the vaccination of bonobos and orangutans with an experimental COVID-19 vaccine [133]. These species may benefit from focused risk mitigation efforts, such as those enacted recently to protect endangered black-footed ferrets (*Mustela nigripes*) from potential SARS-CoV-2 spillback [134].

All fifteen species of *Tupaia* treeshrews were predicted by our model to have medium to high probability (ranging from 0.62 to 0.87). One species, *T. belangeri*, has been explored as a potential lab model for several human infectious diseases including SARS-CoV-2 [135] but relative to other treeshrews, our model assigned only medium probability for SARS-CoV-2 zoonotic capacity in this species (0.67). This result matches lab studies reporting asymptomatic infection and low viral shedding in *T. belangeri* [54]. In contrast, the common treeshrew (*T. glis*) was in the 94th percentile of zoonotic capacity (0.87 probability). These two species are sympatric in parts of their range, exist in close proximity to humans, and also overlap geographically with COVID-19 hotspots in Southeast Asia, suggesting the possibility of spillover transmission among congeners if spillback transmission occurs from humans to these species.

### Strengthening predictive capacity for zoonoses

While there was wide agreement between our model predictions and empirical studies, examining biases and mismatches between experimental results and model-generated predictions will focus research attention on characterizing what factors underlie the disconnects between predicted and observed zoonotic capacity. For instance, this study along with multiple other computational and experimental studies predicted that pigs (*Sus scrofa*) would be susceptible to SARS-CoV-2 (Figure 1), but this prediction has not been supported by results from whole animal inoculations [37,53].

Disconnects between real-world observations, *in vivo* experimental results, and *in silico* predictions of zoonotic capacity may arise because host susceptibility and transmission capacity are necessary but not sufficient for zoonotic risk to be realized in natural settings. These processes are embedded in a broader ecological context that impacts host susceptibility, intra-host infection dynamics (latency, recrudescence, tolerance), and viral persistence that collectively determine where and when spillover will occur [136–139]. These processes also depend strongly on the cellular environments in which cell entry and viral replication take place (e.g., the presence of key proteases, [7]), and on host immunogenicity [139], factors which are themselves influenced by the environment [140]. Insofar as data limitations preclude perfect computational predictions of zoonotic capacity (e.g., limited ACE2 sequences, crystal structures, or species trait data), laboratory experiments are also limited in assessing true zoonotic capacity. For SARS-CoV-2 and other host-pathogen systems, animals that are readily infected in the lab appear to be less susceptible in non-lab settings (ferrets in the lab vs. mixed results in ferrets as pets [36,53,141]; rabbits in the lab vs. rabbits as pets [48,142]. Moreover, wildlife hosts confirmed to shed multiple zoonotic viruses in natural settings (e.g., bats, [143]) can be much less tractable for whole-animal laboratory investigations (for instance, requiring high biosecurity containment and very limited sample sizes in unnatural settings). While laboratory experiments are critical for understanding mechanisms of pathogenesis and disease, without field surveillance and population-level studies they offer imperfect reflections of zoonotic capacity in the natural world.

These examples illustrate that there is no single methodology sufficient to understand and predict zoonotic transmission, for SARS-CoV-2 or any zoonotic pathogen. They also demonstrate the need for improved coordination among theoretical and statistical models, lab work, and field work to improve zoonotic predictive capacity [144], and to create new linkages to underutilized data sources such as natural history collections, which are well-positioned to augment basic knowledge gaps about the spatial and temporal extents of animal hosts and their pathogens [145,146]. Integration of multiple methodologies and data streams across biological scales offers avenues to more efficient iteration between computational predictions, laboratory experiments, and targeted animal surveillance that will better link transmission mechanisms to the broader conditions underpinning zoonotic disease emergence in nature.

## Acknowledgments

We are grateful for discussions with Drs. Alexandre Bonvin, Dennis Bente, Susan Hafenstein, Kathryn Hanley, Hyunwook Lee, Colin Parrish, and John Paul Schmidt about various components of this project. This work was supported by the NSF EEID program (DEB 1717282), DARPA PREEMPT program (D18AC00031), CREATE-NEO, a member of the NIH NIAID CREID program (1U01 AI151807-01), and the NVIDIA Corporation GPU grant program (BAH); by the NSF Polar program (OPP 1935870, 1947040) (AV); and by NIH NIGMS (R35GM122543) (JPGLMR).

## References

1. Dong E, Du H, Gardner L. 2020 An interactive web-based dashboard to track COVID-19 in real time. Lancet Infect. Dis. 20, 533–534. (doi:10.1016/S1473-3099(20)30120-1)

2. WHO. 2021 WHO coronavirus disease (COVID-19) dashboard.

3. Keele BF et al. 2006 Chimpanzee reservoirs of pandemic and nonpandemic HIV-1. Science 313, 523–526. (doi:10.1126/science.1126531)

4. Gage KL, Kosoy MY. 2005 Natural history of plague: perspectives from more than a century of research. Annu. Rev. Entomol. 50, 505–528. (doi:10.1146/annurev.ento.50.071803.130337)

5. Taubenberger JK, Reid AH, Lourens RM, Wang R, Jin G, Fanning TG. 2005 Characterization of the 1918 influenza virus polymerase genes. Nature 437, 889. (doi:10.1038/nature04230)

6. Zhou P et al. 2020 A pneumonia outbreak associated with a new coronavirus of probable bat origin. Nature 579, 270–273. (doi:10.1038/s41586-020-2012-7)

7. Letko M, Marzi A, Munster V. 2020 Functional assessment of cell entry and receptor usage for SARS-CoV-2 and other lineage B betacoronaviruses. Nat Microbiol 5, 562–569. (doi:10.1038/s41564-020-0688-y)

8. Chou C-F et al. 2006 ACE2 orthologues in non-mammalian vertebrates (Danio, Gallus, Fugu, Tetraodon and Xenopus). Gene 377, 46–55. (doi:10.1016/j.gene.2006.03.010)

9. Guth S, Visher E, Boots M, Brook CE. 2019 Host phylogenetic distance drives trends in virus virulence and transmissibility across the animal-human interface. Philos. Trans. R. Soc. Lond. B Biol. Sci. 374, 20190296. (doi:10.1098/rstb.2019.0296)

10. WHO. 2020 SARS-CoV-2 mink-associated variant strain – Denmark.

11. Oude Munnink BB et al. 2020 Transmission of SARS-CoV-2 on mink farms between humans and mink and back to humans. Science (doi:10.1126/science.abe5901)

12. Garry RF. 2021 Mutations arising in SARS-CoV-2 spike on sustained human-to-human transmission and human-to-animal passage. Virological. See https://virological.org/t/mutations-arising-in-sars-cov-2-spike-on-sustained-human-to-human-transmission-and-human-to-animal-passage/578 (accessed on 28 January 2021).

13. Rodrigues JPGLM, Barrera-Vilarmau S, M C Teixeira J, Sorokina M, Seckel E, Kastritis PL, Levitt M. 2020 Insights on cross-species transmission of SARS-CoV-2 from structural modeling. PLoS Comput. Biol. 16, e1008449. (doi:10.1371/journal.pcbi.1008449)

14. Davies NG et al. 2020 Estimated transmissibility and severity of novel SARS-CoV-2 Variant of Concern 202012/01 in England. medRxiv, 2020.12.24.20248822. (doi:10.1101/2020.12.24.20248822)

15. Volz E et al. 2021 Transmission of SARS-CoV-2 Lineage B.1.1.7 in England: Insights from linking epidemiological and genetic data. medRxiv, 2020.12.30.20249034. (doi:10.1101/2020.12.30.20249034)

16. Rambaut A et al. 2020 Preliminary genomic characterisation of an emergent SARS-CoV-2 lineage in the UK defined by a novel set of spike mutations. Virological. See https://virological.org/t/preliminary-genomic-characterisation-of-an-emergent-sars-cov-2-lineage-in-the-uk-defined-by-a-novel-set-of-spike-mutations/563 (accessed on 28 January 2021).

17. Tegally H et al. 2020 Emergence and rapid spread of a new severe acute respiratory syndrome-related coronavirus 2 (SARS-CoV-2) lineage with multiple spike mutations in South Africa. medRxiv, 2020.12.21.20248640. (doi:10.1101/2020.12.21.20248640)

18. Van Egeren D et al. 2021 Risk of rapid evolutionary escape from biomedical interventions targeting SARS-CoV-2 spike protein. PLoS One 16, e0250780. (doi:10.1371/journal.pone.0250780)

19. Bao L et al. 2020 The pathogenicity of SARS-CoV-2 in hACE2 transgenic mice. Nature 583, 830–833. (doi:10.1038/s41586-020-2312-y)

20. Montagutelli X et al. 2021 The B1.351 and P.1 variants extend SARS-CoV-2 host range to mice. bioRxiv., 2021.03.18.436013. (doi:10.1101/2021.03.18.436013)

21. DeLiberto T, Shriner S. 2020 ProMED.

22. ODA. 2020 Mink at affected Oregon farm negative for SARS-CoV-2, wildlife surveillance continues. 23 December. See https://odanews.wpengine.com/mink-at-affected-oregon-farm-negative-for-sars-cov-2-wildlife-surveillance-continues/.

23. Shriner S et al. 2021 SARS-CoV-2 Exposure in Escaped Mink, Utah, USA. Emerging Infectious Disease journal 27. (doi:10.3201/eid2703.204444)

24. Lam SD et al. 2020 SARS-CoV-2 spike protein predicted to form complexes with host receptor protein orthologues from a broad range of mammals. Sci. Rep. 10, 16471. (doi:10.1038/s41598-020-71936-5)

25. Liu Z et al. 2020 Composition and divergence of coronavirus spike proteins and host ACE2 receptors predict potential intermediate hosts of SARS-CoV-2. J. Med. Virol. 92, 595–601. (doi:10.1002/jmv.25726)

26. Luan J, Jin X, Lu Y, Zhang L. 2020 SARS-CoV-2 spike protein favors ACE2 from Bovidae and Cricetidae. J. Med. Virol.

27. Mathavarajah S, Stoddart AK, Gagnon GA, Dellaire G. 2020 Pandemic danger to the deep: the risk of marine mammals contracting SARS-CoV-2 from wastewater., 2020.08.13.249904. (doi:10.1101/2020.08.13.249904)

28. Melin AD, Janiak MC, Marrone F 3rd, Arora PS, Higham JP. 2020 Comparative ACE2 variation and primate COVID-19 risk. Commun Biol 3, 641. (doi:10.1038/s42003-020-01370-w)

29. Kumar A, Pandey SN, Pareek V, Narayan RK, Faiq MA, Kumari C. 2020 Predicting susceptibility for SARS-CoV-2 infection in domestic and wildlife animals using ACE2 protein sequence homology. Zoo Biol. (doi:10.1002/zoo.21576)

30. Huang X, Zhang C, Pearce R, Omenn GS, Zhang Y. 2020 Identifying the Zoonotic Origin of SARS-CoV-2 by Modeling the Binding Affinity between the Spike Receptor-Binding Domain and Host ACE2. J. Proteome Res. 19, 4844–4856. (doi:10.1021/acs.jproteome.0c00717)

31. Ahmed R, Hasan R, Siddiki Amamz, Islam MS. 2021 Host range projection of SARS-CoV-2: South Asia perspective. Infect. Genet. Evol. 87, 104670. (doi:10.1016/j.meegid.2020.104670)

32. Damas J et al. 2020 Broad host range of SARS-CoV-2 predicted by comparative and structural analysis of ACE2 in vertebrates. Proc. Natl. Acad. Sci. U. S. A. 117, 22311– 22322. (doi:10.1073/pnas.2010146117)

33. Ulrich L, Wernike K, Hoffmann D, Mettenleiter TC, Beer M. 2020 Experimental Infection of Cattle with SARS-CoV-2. Emerg. Infect. Dis. 26, 2979–2981. (doi:10.3201/eid2612.203799)

34. Sit THC et al. 2020 Infection of dogs with SARS-CoV-2. Nature 586, 776–778. (doi:10.1038/s41586-020-2334-5)

35. USDA. 2020 Cases of SARS-CoV-2 in animals in the United States. See https://www.aphis.usda.gov/aphis/ourfocus/animalhealth/sa_one_health/sars-cov-2-animals-us (accessed on 12 January 2021).

36. OIE. 2021 Events in animals: OIE - World Organisation for Animal Health. See https://www.oie.int/en/scientific-expertise/specific-information-and-recommendations/questions-and-answers-on-2019novel-coronavirus/events-in-animals/ (accessed on 28 January 2021).

37. Shi J et al. 2020 Susceptibility of ferrets, cats, dogs, and other domesticated animals to SARS-coronavirus 2. Science 368, 1016–1020. (doi:10.1126/science.abb7015)

38. Hamer SA et al. 2021 SARS-CoV-2 Infections and Viral Isolations among Serially Tested Cats and Dogs in Households with Infected Owners in Texas, USA. Viruses 13. (doi:10.3390/v13050938)

39. Woolsey C et al. 2020 Establishment of an African green monkey model for COVID-19. bioRxiv, 2020.05.17.100289. (doi:10.1101/2020.05.17.100289)

40. Hall JS et al. 2020 Experimental challenge of a North American bat species, big brown bat (Eptesicus fuscus), with SARS-CoV-2. Transbound. Emerg. Dis. (doi:10.1111/tbed.13949)

41. Zhang Q et al. 2020 SARS-CoV-2 neutralizing serum antibodies in cats: a serological investigation., 2020.04.01.021196. (doi:10.1101/2020.04.01.021196)

42. San Diego Zoo. 2021 Gorilla Troop at the San Diego Zoo Safari Park Test Positive for COVID-19. See https://zoo.sandiegozoo.org/pressroom/news-releases/gorilla-troop-san-diego-zoo-safari-park-test-positive-covid-19 (accessed on 28 January 2021).

43. Rockx B et al. 2020 Comparative pathogenesis of COVID-19, MERS, and SARS in a nonhuman primate model. Science 368, 1012–1015. (doi:10.1126/science.abb7314)

44. Munster VJ et al. 2020 Respiratory disease in rhesus macaques inoculated with SARS-CoV-2. Nature 585, 268–272. (doi:10.1038/s41586-020-2324-7)

45. Sia SF et al. 2020 Pathogenesis and transmission of SARS-CoV-2 in golden hamsters. Nature 583, 834–838. (doi:10.1038/s41586-020-2342-5)

46. Oreshkova N et al. 2020 SARS-CoV-2 infection in farmed minks, the Netherlands, April and May 2020. Euro Surveill. 25. (doi:10.2807/1560-7917.ES.2020.25.23.2001005)

47. Freuling CM et al. 2020 Susceptibility of Raccoon Dogs for Experimental SARS-CoV-2 Infection. Emerg. Infect. Dis. 26, 2982–2985. (doi:10.3201/eid2612.203733)

48. Mykytyn AZ et al. 2021 Susceptibility of rabbits to SARS-CoV-2. Emerg. Microbes Infect. 10, 1–7. (doi:10.1080/22221751.2020.1868951)

49. Bartlett SL et al. 2021 SARS-COV-2 INFECTION AND LONGITUDINAL FECAL SCREENING IN MALAYAN TIGERS (PANTHERA TIGRIS JACKSONI), AMUR TIGERS (PANTHERA TIGRIS ALTAICA), AND AFRICAN LIONS (PANTHERA LEO KRUGERI) AT THE BRONX ZOO, NEW YORK, USA. J. Zoo Wildl. Med. 51, 733–744. (doi:10.1638/2020-0171)

50. Wang L et al. 2020 Complete Genome Sequence of SARS-CoV-2 in a Tiger from a U.S. Zoological Collection. Microbiol Resour Announc 9. (doi:10.1128/MRA.00468-20)

51. Fagre A et al. 2021 SARS-CoV-2 infection, neuropathogenesis and transmission among deer mice: Implications for spillback to New World rodents. PLoS Pathog. 17, e1009585. (doi:10.1371/journal.ppat.1009585)

52. Griffin BD et al. 2021 SARS-CoV-2 infection and transmission in the North American deer mouse. Nat. Commun. 12, 3612. (doi:10.1038/s41467-021-23848-9)

53. Schlottau K et al. 2020 SARS-CoV-2 in fruit bats, ferrets, pigs, and chickens: an experimental transmission study. Lancet Microbe 1, e218–e225. (doi:10.1016/S2666-5247(20)30089-6)

54. Zhao Y et al. 2020 Susceptibility of tree shrew to SARS-CoV-2 infection. Sci. Rep. 10, 16007. (doi:10.1038/s41598-020-72563-w)

55. Louisville Zoo. 2020 Louisville Zoo Female Snow Leopard Tests Positive for SARS-CoV-2. See https://louisvillezoo.org/louisville-zoo-female-snow-leopard-tests-positive-for-sars-cov-2-media-release/ (accessed on 28 January 2021).

56. Ulrich L, Michelitsch A, Halwe N, Wernike K, Hoffmann D, Beer M. 2021 Experimental SARS-CoV-2 Infection of Bank Voles. Emerg. Infect. Dis. 27, 1193–1195. (doi:10.3201/eid2704.204945)

57. Georgia Aquarium. In press. Asian Small-Clawed Otters at Georgia Aquarium Test Positive for COVID-19. See http://news.georgiaaquarium.org/stories/releases-20210418 (accessed on 13 May 2021).

58. Palmer MV et al. 2021 Susceptibility of white-tailed deer (Odocoileus virginianus) to SARS-CoV-2. bioRxiv., 2021.01.13.426628. (doi:10.1101/2021.01.13.426628)

59. Gryseels S, De Bruyn L, Gyselings R, Calvignac-Spencer S, Leendertz FH, Leirs H. 2020 Risk of human-to-wildlife transmission of SARS-CoV-2. Mamm. Rev. 8, e00373–17. (doi:10.1111/mam.12225)

60. Deng W et al. 2020 Rhesus macaques can be effectively infected with SARS-CoV-2 via ocular conjunctival route. bioRxiv., 2020.03.13.990036. (doi:10.1101/2020.03.13.990036)

61. Rodrigues Jpglm et al. 2013 Defining the limits of homology modeling in information-driven protein docking. Proteins 81, 2119–2128. (doi:10.1002/prot.24382)

62. Sander C, Schneider R. 1991 Database of homology-derived protein structures and the structural meaning of sequence alignment. Proteins 9, 56–68. (doi:10.1002/prot.340090107)

63. Li Y et al. 2020 SARS-CoV-2 and Three Related Coronaviruses Utilize Multiple ACE2 Orthologs and Are Potently Blocked by an Improved ACE2-Ig. J. Virol. 94. (doi:10.1128/JVI.01283-20)

64. Fournier D, Luft FC, Bader M, Ganten D, Andrade-Navarro MA. 2012 Emergence and evolution of the renin-angiotensin-aldosterone system. J. Mol. Med. 90, 495–508. (doi:10.1007/s00109-012-0894-z)

65. Han BA, Schmidt JP, Bowden SE, Drake JM. 2015 Rodent reservoirs of future zoonotic diseases. Proc. Natl. Acad. Sci. U. S. A. 112, 7039–7044. (doi:10.1073/pnas.1501598112)

66. Yang LH, Han BA. 2018 Data-driven predictions and novel hypotheses about zoonotic tick vectors from the genus Ixodes. BMC Ecol. 18, 7. (doi:10.1186/s12898-018-0163-2)

67. Han BA, O’Regan SM, Paul Schmidt J, Drake JM. 2020 Integrating data mining and transmission theory in the ecology of infectious diseases. Ecol. Lett. 23, 1178–1188. (doi:10.1111/ele.13520)

68. Han BA, Schmidt JP, Alexander LW, Bowden SE, Hayman DTS, Drake JM. 2016 Undiscovered Bat Hosts of Filoviruses. PLoS Negl. Trop. Dis. 10, e0004815. (doi:10.1371/journal.pntd.0004815)

69. Han BA et al. 2019 Confronting data sparsity to identify potential sources of Zika virus spillover infection among primates. Epidemics 27, 59–65. (doi:10.1016/j.epidem.2019.01.005)

70. Yang X-L et al. 2017 Genetically Diverse Filoviruses in Rousettus and Eonycteris spp. Bats, China, 2009 and 2015. Emerg. Infect. Dis. 23, 482–486. (doi:10.3201/eid2303.161119)

71. Goldstein T et al. 2018 The discovery of Bombali virus adds further support for bats as hosts of ebolaviruses. Nat Microbiol 3, 1084–1089. (doi:10.1038/s41564-018-0227-2)

72. Sorokina M, M C Teixeira J, Barrera-Vilarmau S, Paschke R, Papasotiriou I, Rodrigues Jpglm, Kastritis PL. 2020 Structural models of human ACE2 variants with SARS-CoV-2 Spike protein for structure-based drug design. Sci Data 7, 309. (doi:10.1038/s41597-020-00652-6)

73. Altschul SF, Gish W, Miller W, Myers EW, Lipman DJ. 1990 Basic local alignment search tool. J. Mol. Biol. 215, 403–410. (doi:10.1016/S0022-2836(05)80360-2)

74. Winter D. 2017 rentrez: An R package for the NCBI eUtils API. R J. 9, 520. (doi:10.32614/rj-2017-058)

75. Rawlings ND, Barrett AJ, Thomas PD, Huang X, Bateman A, Finn RD. 2018 The MEROPS database of proteolytic enzymes, their substrates and inhibitors in 2017 and a comparison with peptidases in the PANTHER database. Nucleic Acids Res. 46, D624–D632. (doi:10.1093/nar/gkx1134)

76. Katoh K, Misawa K, Kuma K-I, Miyata T. 2002 MAFFT: a novel method for rapid multiple sequence alignment based on fast Fourier transform. Nucleic Acids Res. 30, 3059–3066. (doi:10.1093/nar/gkf436)

77. Lan J et al. 2020 Structure of the SARS-CoV-2 spike receptor-binding domain bound to the ACE2 receptor. Nature 581, 215–220. (doi:10.1038/s41586-020-2180-5)

78. Webb B, Sali A. 2016 Comparative Protein Structure Modeling Using MODELLER. Curr. Protoc. Bioinformatics 54, 5.6.1–5.6.37. (doi:10.1002/cpbi.3)

79. Sali A, Blundell TL. 1993 Comparative protein modelling by satisfaction of spatial restraints. J. Mol. Biol. 234, 779–815. (doi:10.1006/jmbi.1993.1626)

80. van Zundert GCP et al. 2016 The HADDOCK2.2 Web Server: User-Friendly Integrative Modeling of Biomolecular Complexes. J. Mol. Biol. 428, 720–725. (doi:10.1016/j.jmb.2015.09.014)

81. de Magalhães JP, Costa J. 2009 A database of vertebrate longevity records and their relation to other life-history traits. J. Evol. Biol. 22, 1770–1774. (doi:10.1111/j.1420-9101.2009.01783.x)

82. Myhrvold NP, Baldridge E, Chan B, Sivam D, Freeman DL, Ernest SKM. 2015 An amniote life-history database to perform comparative analyses with birds, mammals, and reptiles: Ecological ArchivesE096-269. Ecology 96, 3109–3000. (doi:10.1890/15-0846r.1)

83. Wilman H, Belmaker J, Simpson J, de la Rosa C, Rivadeneira MM, Jetz W. 2014 EltonTraits 1.0: Species-level foraging attributes of the world’s birds and mammals. Ecology 95, 2027–2027. (doi:10.1890/13-1917.1)

84. Dallas T, Park AW, Drake JM. 2017 Predicting cryptic links in host-parasite networks. PLoS Comput. Biol. 13, e1005557. (doi:10.1371/journal.pcbi.1005557)

85. Baker C. 2018 wosr: Clients to the ‘Web of Science’ and ‘InCites’ APIs. See https://CRAN.R-project.org/package=wosr.

86. Bosco-Lauth AM et al. 2020 Experimental infection of domestic dogs and cats with SARS-CoV-2: Pathogenesis, transmission, and response to reexposure in cats. Proc. Natl. Acad. Sci. U. S. A. 117, 26382–26388. (doi:10.1073/pnas.2013102117)

87. Elith J, Leathwick JR, Hastie T. 2008 A working guide to boosted regression trees. J. Anim. Ecol. 77, 802–813.

88. Jones KE et al. 2009 PanTHERIA: a species-level database of life history, ecology, and geography of extant and recently extinct mammals: Ecological Archives E090-184. Ecology 90, 2648–2648. (doi:10.1890/08-1494.1)

89. Wilson DE, Reeder DM. 2005 Mammal Species of the World: A Taxonomic and Geographic Reference. Baltimore, MD: JHU Press. See http://books.google.com/books?id=JgAMbNSt8ikC.

90. Greenwell B, Boehmke B, Cunningham J, Developers GBM. 2020 Generalized Boosted Regression Models. Comprehensive R Archive Network (CRAN). See https://cran.r-project.org/web/packages/gbm/index.html.

91. R Core Team. 2020 R: A language and environment for statistical computing. Vienna, Austria. See http://www.R-project.org/.

92. Wilkins AS, Wrangham RW, Fitch WT. 2014 The ‘domestication syndrome’ in mammals: a unified explanation based on neural crest cell behavior and genetics. Genetics 197, 795– 808. (doi:10.1534/genetics.114.165423)

93. Cleaveland S, Laurenson MK, Taylor LH. 2001 Diseases of humans and their domestic mammals: pathogen characteristics, host range and the risk of emergence. Philos. Trans. R. Soc. Lond. B Biol. Sci. 356, 991–999. (doi:10.1098/rstb.2001.0889)

94. Database MD. 2020 Mammal Diversity Database. (doi:10.5281/zenodo.4139818)

95. IUCN. 2020 The IUCN Red List of Threatened Species.

96. Liu Y et al. 2021 Functional and genetic analysis of viral receptor ACE2 orthologs reveals a broad potential host range of SARS-CoV-2. Proc. Natl. Acad. Sci. U. S. A. 118. (doi:10.1073/pnas.2025373118)

97. Pitra C, Schwarz S, Fickel J. 2010 Going west—invasion genetics of the alien raccoon dog Nyctereutes procynoides in Europe. Eur. J. Wildl. Res. 56, 117–129. (doi:10.1007/s10344-009-0283-2)

98. Milla R et al. 2018 Phylogenetic patterns and phenotypic profiles of the species of plants and mammals farmed for food. Nat Ecol Evol 2, 1808–1817. (doi:10.1038/s41559-018-0690-4)

99. Kevany S. 2020 Danish Covid mink cull and future disease fears will kill fur trade, say farmers. The Guardian, 6 November. See http://www.theguardian.com/environment/2020/nov/06/danish-covid-mink-cull-and-future-disease-fears-will-kill-fur-trade-say-farmers.

100. Can ÖE, D’Cruze N, Macdonald DW. 2019 Dealing in deadly pathogens: Taking stock of the legal trade in live wildlife and potential risks to human health. Glob Ecol Conserv 17, e00515. (doi:10.1016/j.gecco.2018.e00515)

101. Lam TT-Y et al. 2020 Identifying SARS-CoV-2-related coronaviruses in Malayan pangolins. Nature 583, 282–285. (doi:10.1038/s41586-020-2169-0)

102. Andersen KG, Rambaut A, Lipkin WI, Holmes EC, Garry RF. 2020 The proximal origin of SARS-CoV-2. Nat. Med. 26, 450–452. (doi:10.1038/s41591-020-0820-9)

103. Lehmann D, Halbwax ML, Makaga L, Whytock R, Ndindiwe Malata L, Bombenda Mouele W, Momboua BR, Koumba Pambo AF, White LJT. 2020 Pangolins and bats living together in underground burrows in Lopé National Park, Gabon. Afr. J. Ecol. (doi:10.1111/aje.12759)

104. Tsuda S et al. 2012 Genomic and serological detection of bat coronavirus from bats in the Philippines. Arch. Virol. 157, 2349–2355. (doi:10.1007/s00705-012-1410-z)

105. Olival KJ et al. 2020 Possibility for reverse zoonotic transmission of SARS-CoV-2 to free-ranging wildlife: A case study of bats. PLoS Pathog. 16, e1008758. (doi:10.1371/journal.ppat.1008758)

106. Anthony SJ et al. 2013 Coronaviruses in bats from Mexico. J. Gen. Virol. 94, 1028–1038. (doi:10.1099/vir.0.049759-0)

107. Anthony SJ et al. 2017 Global patterns in coronavirus diversity. Virus Evol 3, vex012. (doi:10.1093/ve/vex012)

108. Zhou H et al. 2020 A Novel Bat Coronavirus Closely Related to SARS-CoV-2 Contains Natural Insertions at the S1/S2 Cleavage Site of the Spike Protein. Curr. Biol. 30, 2196–2203.e3. (doi:10.1016/j.cub.2020.05.023)

109. Hul V et al. 2021 A novel SARS-CoV-2 related coronavirus in bats from Cambodia. bioRxiv., 2021.01.26.428212. (doi:10.1101/2021.01.26.428212)

110. Mou H et al. 2020 Mutations from bat ACE2 orthologs markedly enhance ACE2-Fc neutralization of SARS-CoV-2. bioRxiv., 2020.06.29.178459. (doi:10.1101/2020.06.29.178459)

111. Wacharapluesadee S et al. 2021 Evidence for SARS-CoV-2 related coronaviruses circulating in bats and pangolins in Southeast Asia. Nat. Commun. 12, 972. (doi:10.1038/s41467-021-21240-1)

112. Pulliam JRC et al. 2012 Agricultural intensification, priming for persistence and the emergence of Nipah virus: a lethal bat-borne zoonosis. J. R. Soc. Interface 9, 89–101. (doi:10.1098/rsif.2011.0223)

113. Plowright RK et al. 2015 Ecological dynamics of emerging bat virus spillover. Proceedings of the Royal Society B. 282, 20142124. (doi:10.1098/rspb.2014.2124)

114. Kessler MK et al. 2018 Changing resource landscapes and spillover of henipaviruses. Ann. N. Y. Acad. Sci. 1429, 78–99. (doi:10.1111/nyas.13910)

115. Peel AJ, Field HE, Aravena MR, Edson D, McCallum H, Plowright RK, Prada D. 2020 Coronaviruses and Australian bats: a review in the midst of a pandemic. Aust. J. Zool. (doi:10.1071/ZO20046)

116. Bordes F, Blasdell K, Morand S. 2015 Transmission ecology of rodent-borne diseases: New frontiers. Integr. Zool. 10, 424–435. (doi:10.1111/1749-4877.12149)

117. Ostfeld RS, Canham CD, Oggenfuss K, Winchcombe RJ, Keesing F. 2006 Climate, deer, rodents, and acorns as determinants of variation in lyme-disease risk. PLoS Biol. 4, e145. (doi:10.1371/journal.pbio.0040145)

118. Machtinger ET, Williams SC. 2020 Practical Guide to Trapping Peromyscus leucopus (Rodentia: Cricetidae) and Peromyscus maniculatus for Vector and Vector-Borne Pathogen Surveillance and Ecology. J. Insect Sci. 20. (doi:10.1093/jisesa/ieaa028)

119. Morand S et al. 2015 Global parasite and Rattus rodent invasions: The consequences for rodent-borne diseases. Integr. Zool. 10, 409–423. (doi:10.1111/1749-4877.12143)

120. Hamdan NES, Ng YL, Lee WB, Tan CS, Khan FAA, Chong YL. 2017 Rodent Species Distribution and Hantavirus Seroprevalence in Residential and Forested areas of Sarawak, Malaysia. Trop Life Sci Res 28, 151–159. (doi:10.21315/tlsr2017.28.1.11)

121. Louys J, Herrera MB, Thomson VA, Wiewel AS, Donnellan SC, O’Connor S, Aplin K. 2020 Expanding population edge craniometrics and genetics provide insights into dispersal of commensal rats through Nusa Tenggara, Indonesia. Rec. Aust. Mus. 72, 287–302. (doi:10.3853/j.2201-4349.72.2020.1730)

122. Tadin A et al. 2016 Molecular Survey of Zoonotic Agents in Rodents and Other Small Mammals in Croatia. Am. J. Trop. Med. Hyg. 94, 466–473. (doi:10.4269/ajtmh.15-0517)

123. Yousefi A, Eslami A, Mobedi I, Rahbari S, Ronaghi H. 2014 Helminth Infections of House Mouse (Mus musulus) and Wood Mouse (Apodemus sylvaticus) from the Suburban Areas of Hamadan City, Western Iran. Iran. J. Parasitol. 9, 511–518.

124. Rahman MM, Yoon KB, Lim SJ, Jeon MG, Kim HJ, Kim HY, Cho JY, Chae HM, Park YC. 2017 Molecular detection by analysis of the 16S rRNA gene of fecal coliform bacteria from the two Korean Apodemus species (Apodemus agrarius and A. peninsulae). Genet. Mol. Res. 16. (doi:10.4238/gmr16029510)

125. Bertolino S, Lurz PWW. 2013 Callosciurussquirrels: worldwide introductions, ecological impacts and recommendations to prevent the establishment of new invasive populations: Worldwide introductions ofCallosciurussquirrels. Mamm. Rev. 43, 22–33. (doi:10.1111/j.1365-2907.2011.00204.x)

126. Mildenstein T, Tanshi I, Racey PA. 2016 Exploitation of Bats for Bushmeat and Medicine. In Bats in the Anthropocene: Conservation of Bats in a Changing World (eds CC Voigt, T Kingston), pp. 325–375. Cham: Springer International Publishing. (doi:10.1007/978-3-319-25220-9_12)

127. Ransaleleh TA, Nangoy MJ, Wahyuni I, Lomboan A, Koneri R, Saputro S, Pamungkas J, Latinne A. 2020 Identification of bats on traditional market in dumoga district, North Sulawesi. IOP Conf. Ser.: Earth Environ. Sci. 473, 012067. (doi:10.1088/1755-1315/473/1/012067)

128. Tampon NVT, Rabaya YMC, Malbog KMA, Burgos SC, Libre K Jr, Valila ASD, Achondo Mjmm, Onggo LS, Murao LAE. 2020 First molecular evidence for bat betacoronaviruses in Mindanao. Philipp J. Sci. 149, 91–94.

129. Logeot M, Mauroy A, Thiry E, De Regge N, Vervaeke M, Beck O, De Waele V, Van den Berg T. 2021 Risk assessment of SARS-CoV-2 infection in free-ranging wild animals in Belgium. Transbound. Emerg. Dis. (doi:10.1111/tbed.14131)

130. Weber A, Kalema-Zikusoka G, Stevens NJ. 2020 Lack of Rule-Adherence During Mountain Gorilla Tourism Encounters in Bwindi Impenetrable National Park, Uganda, Places Gorillas at Risk From Human Disease. Front Public Health 8, 1. (doi:10.3389/fpubh.2020.00001)

131. Barbosa A et al. 2021 Risk assessment of SARS-CoV-2 in Antarctic wildlife. Sci. Total Environ. 755, 143352. (doi:10.1016/j.scitotenv.2020.143352)

132. Gibbons A. 2021 Captive gorillas test positive for coronavirus. Science (doi:10.1126/science.abg5458)

133. Daly N. 2021 First great apes at U.S. zoo receive COVID-19 vaccine made for animals. National Geographic See https://www.nationalgeographic.com/animals/article/first-great-apes-at-us-zoo-receive-coronavirus-vaccine-made-for-animals.

134. Aleccia J. 2020 ‘The Biggest Nemesis’: Black-Footed Ferrets Get Experimental Coronavirus Vaccine. Kaiser Health News, 27 December. See https://www.cpr.org/2020/12/27/the-biggest-nemesis-black-footed-ferrets-get-experimental-coronavirus-vaccine/.

135. Xu L et al. 2020 COVID-19-like symptoms observed in Chinese tree shrews infected with SARS-CoV-2. Zool Res 41, 517–526. (doi:10.24272/j.issn.2095-8137.2020.053)

136. Becker DJ, Washburne AD, Faust CL, Pulliam JRC, Mordecai EA, Lloyd-Smith JO, Plowright RK. 2019 Dynamic and integrative approaches to understanding pathogen spillover. Philos. Trans. R. Soc. Lond. B Biol. Sci. 374, 20190014. (doi:10.1098/rstb.2019.0014)

137. Plowright RK, Parrish CR, McCallum H, Hudson PJ, Ko AI, Graham AL, Lloyd-Smith JO. 2017 Pathways to zoonotic spillover. Nat. Rev. Microbiol. (doi:10.1038/nrmicro.2017.45)

138. Morris DH et al. 2020 The effect of temperature and humidity on the stability of SARS-CoV-2 and other enveloped viruses. bioRxiv (doi:10.1101/2020.10.16.341883)

139. Bean AGD, Baker ML, Stewart CR, Cowled C, Deffrasnes C, Wang L-F, Lowenthal JW. 2013 Studying immunity to zoonotic diseases in the natural host - keeping it real. Nat. Rev. Immunol. 13, 851–861. (doi:10.1038/nri3551)

140. Letko M, Seifert SN, Olival KJ, Plowright RK, Munster VJ. 2020 Bat-borne virus diversity, spillover and emergence. Nat. Rev. Microbiol. 18, 461–471. (doi:10.1038/s41579-020-0394-z)

141. Sawatzki K, Hill NJ, Puryear WB, Foss AD, Stone JJ, Runstadler JA. 2021 Host barriers to SARS-CoV-2 demonstrated by ferrets in a high-exposure domestic setting. Proc. Natl. Acad. Sci. U. S. A. 118. (doi:10.1073/pnas.2025601118)

142. Ruiz-Arrondo I, Portillo A, Palomar AM, Santibanez S, Santibanez P, Cervera C, Oteo JA. 2020 Detection of SARS-CoV-2 in pets living with COVID-19 owners diagnosed during the COVID-19 lockdown in Spain: A case of an asymptomatic cat with SARS-CoV-2 in Europe. bioRxiv. (doi:10.1101/2020.05.14.20101444)

143. Peel AJ et al. 2019 Synchronous shedding of multiple bat paramyxoviruses coincides with peak periods of Hendra virus spillover. Emerg. Microbes Infect. 8, 1314–1323. (doi:10.1080/22221751.2019.1661217)

144. Restif O et al. 2012 Model-guided fieldwork: practical guidelines for multidisciplinary research on wildlife ecological and epidemiological dynamics. Ecol. Lett. (doi:10.1111/j.1461-0248.2012.01836.x)

145. Cook JA et al. 2020 Integrating Biodiversity Infrastructure into Pathogen Discovery and Mitigation of Emerging Infectious Diseases. Bioscience 70, 531–534. (doi:10.1093/biosci/biaa064)

146. Thompson CW et al. 2021 Preserve a Voucher Specimen! The Critical Need for Integrating Natural History Collections in Infectious Disease Studies. MBio 12. (doi:10.1128/mBio.02698-20)

